# Conditions under which distributions of edge length ratios on phylogenetic trees can be used to order evolutionary events

**DOI:** 10.1101/2021.01.16.426961

**Authors:** Edward Susko, Mike Steel, Andrew J. Roger

## Abstract

Two recent high profile studies have attempted to use edge (branch) length ratios from large sets of phylogenetic trees to determine the relative ages of genes of different origins in the evolution of eukaryotic cells. This approach can be straightforwardly justified if substitution rates are constant over the tree for a given protein. However, such strict molecular clock assumptions are not expected to hold on the billion-year timescale. Here we propose an alternative set of conditions under which comparisons of edge length distributions from multiple sets of phylogenies of proteins with different origins can be validly used to discern the order of their origins. We also point out scenarios where these conditions are not expected to hold and caution is warranted.

## Main

The origin of eukaryotic cells from prokaryotic precursors - eukaryogenesis - remains one of the more mysterious major evolutionary transitions in the history of life on Earth. This transition involved a host cellular lineage related to asgard Archaea (Eme *et al.* 2017) that, at some point prior to the last eukaryotic common ancestor (LECA), took up an endosymbiotic alphaproteobacterium that became the mitochondrion, an integrated energy-producing organelle within eukaryotic cells (Dacks *et al.* 2016; Roger *et al.* 2017; Porter 2020). Genes in LECA, therefore, have multiple possible origins: either they were inherited from the host lineage, acquired from the mitochondrial symbiont by endosymbiotic gene transfer, transferred from potentially many other prokaryotic donors by lateral gene transfer (Rochette *et al.* 2014; Pittis and Gabaldón 2016a), or arose *de novo* during eukaryogenesis. Regardless of their origin, many genes were extensively duplicated during this period, as many new cellular traits including the cytoskeleton, nucleus, endomembrane system evolved in the proto-eukaryote lineage. Determining the order of these events remains a major roadblock in our understanding of eukaryogenesis.

In 2016, Pittis and Gabaldón introduced a novel approach to approximating the relative ages of genes of different origins that were acquired during eukaryogenesis (Pittis and Ga-baldón 2016a). The approach relies on the notion that edge lengths on phylogenetic trees estimated from aligned genes or proteins, represent expected numbers of amino acid substitutions along the edge and are proportional to the product of rates of substitution along the edge and the time span of the edge. Under a strict molecular clock assumption the relative lengths of edges are proportional to time spans of the edges.

To characterize the relative timespan a gene has been resident in the proto-eukaryote lineage prior to LECA, Pittis and Gabaldón focused on the edge in each gene tree between the LECA node and the node representing the common ancestor of the closest prokaryotic sister group and the eukaryote lineage (Fig. 1), an edge they call the stem the length of which is denoted here as *L_s_*. All genes from the same origin, *O*, are expected to have a stem edge that corresponds to the same time span (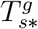 is constant for *g* ∈ *O*). A serious complication arises here almost immediately. For genes in the proto-eukaryote genome that were inherited from its common ancestor with the closest sampled asgard archaeon, the time span of the stem edge in the protein tree 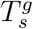 is the same as the timespan it has been resident during eukaryogenesis, 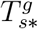. However, for genes that were laterally acquired during eukaryogenesis either *via* the mitochondrial symbiont or from other prokaryotic sources, 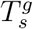 is expected to be larger than 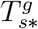 (Fig. 1). This is because the sampled taxa are unlikely to include representatives from the actual immediate prokaryotic sister group of the donor lineage of the gene(s). Reasons for this include inadequacy in sampling of living prokaryote lineages but, more likely, it is because the actual sister group, as opposed to the sampled one, went extinct. In what follows, we assume that for all comparisons 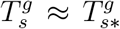, but it is important to recognize the caveats accompanying conclusions coming from stem-length methods applied to comparisons amongst acquired genes. For instance, a claim that the time of acquisition of a group *B* is earlier than that of an acquired group *A*, is more directly an inference that the closest sampled sister lineage of group *B* diverged earlier than that of group *A*.

**Figure 1:**
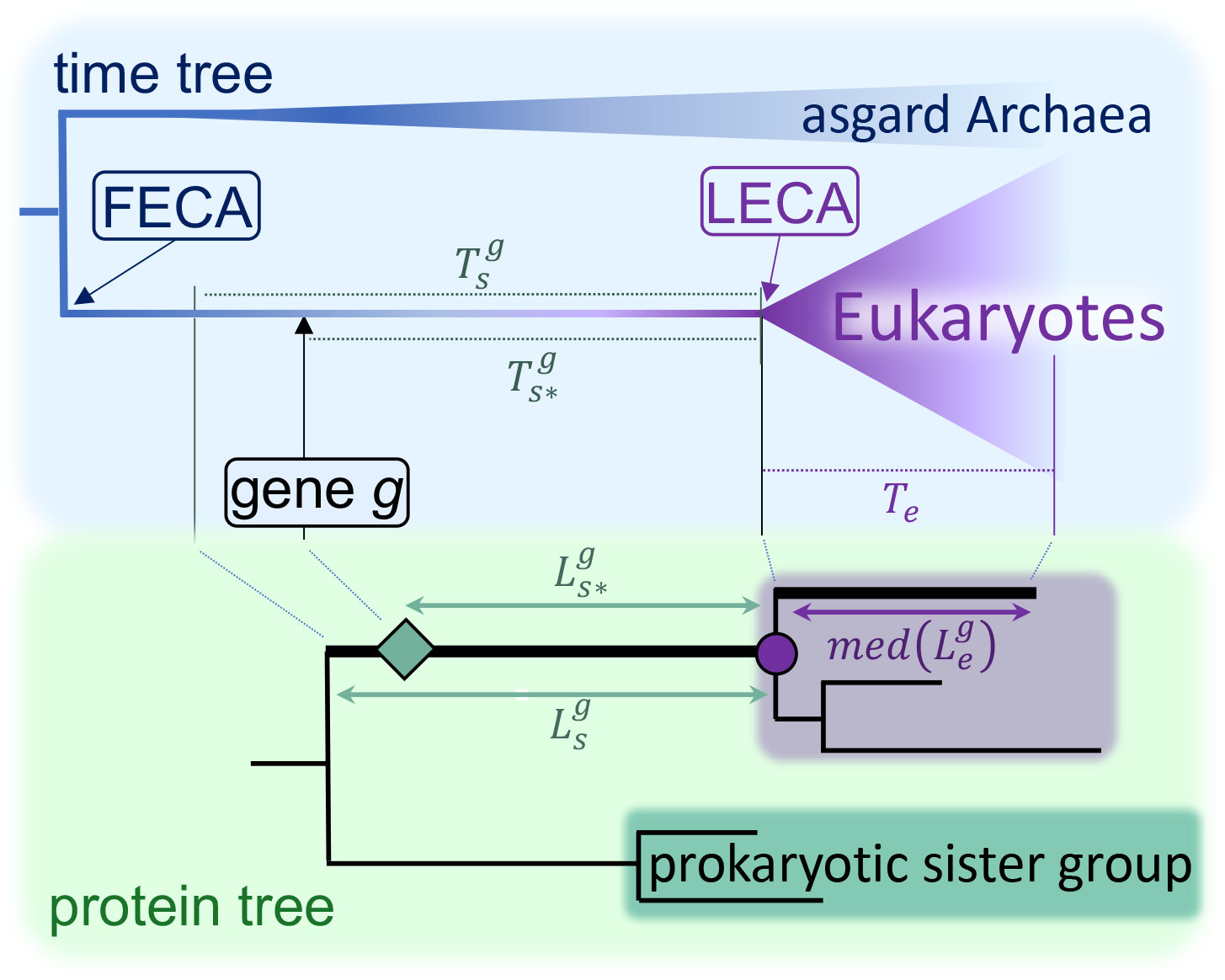
The correspondence between edges on the phylogeny of a gene acquired during eukaryogenesis and edges on the geological time tree of life. The top tree (blue box) shows the part of the geological tree of life depicting the closest asgard archaeal sister group relationship with the ‘host’ lineage of eukaryotes. FECA represents the *first eukaryotic common ancestor* and LECA is the *last eukaryotic common ancestor*. At some point along the eukaryogenesis edge between FECA and LECA, gene *g* was acquired by the proto-eukaryote genome from a prokaryotic lineage. The timespan that gene *g* was present during eukaryogenesis was 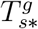 and from LECA to the present is *T_e_*. Below (green box) is the estimated phylogeny of protein *g* and its orthologs. The length of the stem edge is 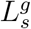 and corresponds to timespan 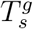 in the time tree. The length of the segment of the latter edge post-acquisition by the proto-eukaryote lineage (green diamond to purple circle) is 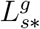 and corresponds to timespan 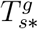. Note that 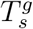, is an upper bound on the timespan of the desired stem edge post-acquisition, 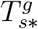, because any gene transfer to the proto-eukaryote from a prokaryotic source must have occurred after speciation of the donor lineage from its closest sampled extant sister group. This discrepancy between the time of origin of a gene and timespan of its stem edge only occurs for genes acquired during eukaryogenesis (e.g. mitochondrial and other laterally acquired genes). The median length of all possible paths between the LECA node (purple circle) and eukaryote leaf node is med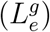. The normalized stem length, 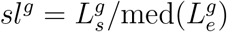.

To remove the effect of different overall rates of evolution in different genes, *R_g_*, Pittis and Gabaldón normalize the original (raw) stem edge length, by a path length, *L_e_*, for a path, *e* (*e* for eukaryote), that corresponds to a constant time span over genes, denoted *T_e_*, giving the time from LECA to the present (Fig. 1). This path length is the sum of consecutive edge lengths over all edges *j* in the path, *L_e_* = Σ*j*∈*e L_j_*. Because for a given protein tree there are multiple paths from LECA to the present, and to exclude potential outliers, the path of median length was chosen as the normalization factor so that the normalized stem length is: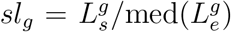. Following Pittis and Gabaldón, we refer to *sl_g_* as a ‘stem length’ even though it is actually a normalized edge length.

Under the molecular clock model, for any path *p*, *L_p_* = *R_g_T_p_*, where *T_p_* is the accumulated time for the path and *R_g_* is the rate of substitution that is constant over time but may vary across genes. Thus, under the molecular clock model, the stem length satisfies that

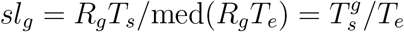

Pittis and Gabaldón (2016a) compared the distributions of estimated stem lengths, 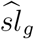 for proteins of different origins. They found that 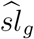 distributions from archaeal origin proteins (*g* ∈ *R*) were, based on Mann-Whitney U tests, significantly shifted to be larger than those of bacterial (*g* ∈ *C*) origin which were, in turn, significantly greater than alphaproteobacterial proteins (*g* ∈ *M*) (the latter are assumed to correspond to genes that originated with the mitochondrial symbiont). Since the times are constant within groups (for instance, 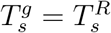 for *g* ∈ *R*), they interpret this as evidence that 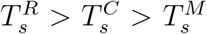. An important conclusion of their study was, therefore, that the mitochondrial symbiosis took place much later in eukaryogenesis than suggested in mitochondria early hypotheses (e.g., Lane and Martin 2010). More recently, Vosseberg and colleagues (Vosseberg *et al.* 2020) have extended this approach to address the relative timings of the mitochondrial symbiosis and gene duplication events for a variety of functional classes of protein families that expanded during eukaryogenesis.

Pittis and Gabaldón’s approach was strongly criticized by Martin and colleagues (Martin *et al.* 2017) who argued that the results were meaningless because the method depends on the assumption that a molecular clock should hold over evolutionary time spans on the billion-year time scale. They investigated a number of the individual phylogenies from Pittis and Gabaldón study and showed that variation in edge lengths within gene trees were substantial and not consistent with a molecular clock. Pittis and Gabaldón have since countered by arguing that their approach does not assume a molecular clock and demonstrate its ability to successfully recover correct orderings of more recent evolutionary divergences in eukaryotes (Pittis and Gabaldón 2016b). However, they did not provide a detailed theoretical justification for why the method should give reliable evolutionary orderings in the absence of a molecular clock

Here we show that assuming a molecular clock is not necessary for the method to work. In what follows, we will show that if proteins *a* and *b* in two groups *a* ∈ *A* and *b* ∈ *B* of different origins evolve independently according to the same time-dependent stochastic substitution rate process, the distributions of the normalized stem lengths of phylogenies of proteins *B* will be systematically larger than *A*, i.e. ℙ[*sl_b_* − *sl_a_* > 0] > 1/2, if and only if 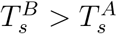. We show that restrictions on the stochastic substitution rate process and the data set required for the foregoing result to hold are surprisingly few, but do include a requirement that there are no systematic differences between groups *A* and *B* at any given time point. This result is shown to hold even when estimation of edge lengths is taken into account, although a modified version of the Mann-Whitney U test will be required for testing when the variances in edge length estimates are systematically different between the groups. We also show that these methods will work in cases in which the proteins within the groups being compared have ranges of different ages. In the latter cases, however, we suggest that statistical test rejection is difficult to meaningfully interpret. Finally, we outline scenarios in which the required assumptions of the method will not hold and caution is warranted.

Borrowing on relaxed molecular clock theory that assumes that the rate of substitution varies stochastically over the tree (Bromham *et al.* 2019), suppose that the rate of substitution at any point along a path *p* can be represented as 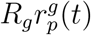, where 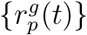 is a continuous time stochastic process (*R_g_* is the overall fixed rate of gene *g* as before). Assuming a conventional Markov substitution model, the probability of substituting state *i* with state *j* in (*t, t*+*h*] is *q_ij_R_g_r_p_*(*t*)*h*+*o*(*h*), for some state transition rate *q_ij_*. Some constraint is required to identify parameters and we assume, without loss of generality, that *E*[*R_g_*] = 1 as well as the conventional constraint, Σ_*i*_ Σ_*j*≠*i*_ *π_i_q_ij_* = 1. Since the chance of two or more substitutions is small relative to *h*, *o*(*h*), the expected number of substitutions in (*t*; *t*+*h*], *E*[*N*(*t*; *t*+*h*)]; is Σ_*i*_ Σ_*j*≠*i*_ *π_i_q_ij_R_g_r_p_*(*t*)*h* + *o*(*h*) = *R_g_r_p_*(*t*)*h* + *o*(*h*). Taking *u*_0_ = 0, *u*_1_ = *t*_0_/*N*,…,*u_N_* = *t*_0_, the number of substitutions in the time period (0; *t*_0_] is the sum, 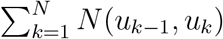 of the substitutions over the intervals (0, *u*_1_],…,(*u_N_*_−1_, *u_N_*]. Since this is true for any *N*, the expected number of substitutions along path *p* and over time period (0*, t*_0_] is

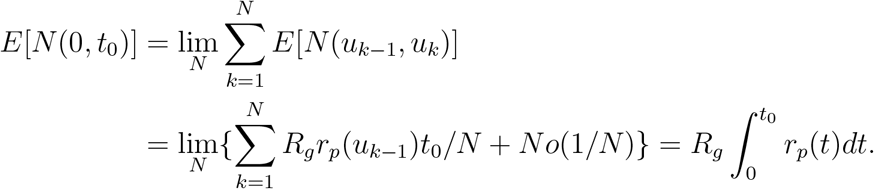

Thus the stem length for protein *g* is

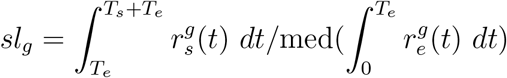

Suppose that comparison is between group *A* and *B* and that 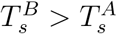. Let

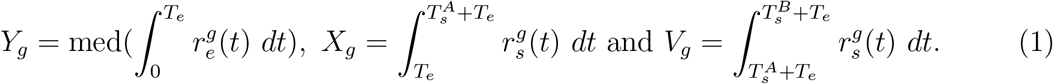

Then *sl_g_* = *X_g_/Y_g_* for *g* ∈ *A* and *sl_g_* = (*X_g_* + *V_g_*)*/Y_g_* for *g* ∈ *B*.

We now make the assumption that for any given path *p* and any two proteins *g* and *g*′ in *A* ∪ *B*, the rate processes are probabilistically equivalent on [0, 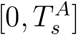]:

For any 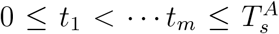 the joint probability distribution of 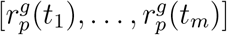 and 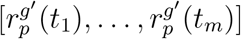 are the same.

Note that this assumption allows possibly radical rate changes throughout the tree and across proteins. Moreover, the rate processes need not be stationary nor Markov processes. What is required, however, is that there be no systematic differences between the two groups. Thus, for instance, the model allows that as a consequence of a radical environmental change or change in population size at time *t*, for some particular lineage *l*, the distribution of 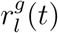 is skewed to the right of the distribution of the rate 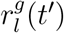 at some other time *t*′. But that difference in distributions is expected to apply whether *g* ∈ *A* or *g* ∈ *B*. For simplicity, we also make the assumption that that the rate process is bounded such that 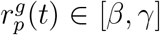 for some *β* > 0 and *γ* < ∞. This assumption is reasonable as we do not expect substitution rates to go to 0 or to increase without bound. In any case, this assumption can be loosened but some sort of assumption is required to avoid having a stem lengths that are almost 0 due entirely to having extremely low average rates along the stem or extremely high average rates from LECA to present. Finally, we assume that the rate processes are independent over genes.

With the assumptions above, if the eukaryotic taxa sampled are the same for groups *A* and *B*, then *Y_g_* will have the same distribution whether *g* ∈ *A* or *g* ∈ *B*. Note that in Pittis and Gabaldón the same taxa were not necessarily present in any two groups of proteins being compared. We argue below that, for the approach to work, it is best if *Y_g_* has the same distribution for *g* ∈ *A* or *g* ∈ *B*, otherwise differences in normalized stem lengths between the groups may be due to unusual rates for eukaryote taxa present in one group but not the other. Thus, it may be desirable to take means or medians over the set of eukaryote taxa present in both groups. Nevertheless, medians are not as likely to be affected by outlying rates (which was one of the original motivations for using them), so if taxon sampling is comparable for the two groups, the distributions of *Y_g_* are expected to be approximately the same for the two groups under the assumptions above.

Consider now ℙ[*sl_b_* − *sl_a_* > *u*] for fixed *u*, *a* ∈ *A* and *b* ∈ *B*. In terms of the random variables above this can be expressed as ℙ[*U* + *V_b_/Y_b_* > *u*] where *U* = *X_b_/Y_b_* − *X_a_/Y_a_*. Since 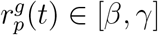, the smallest *V_b_*/*Y_b_* could be is 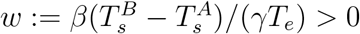. Thus

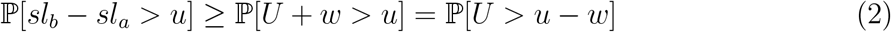

Similarly

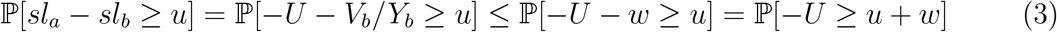

With the assumptions above *X_b_/Y_b_* and *X_a_/Y_a_* have the same distribution. Thus *U* = *X_b_/Y_b_* − *X_a_/Y_a_* has a symmetric distribution around 0. Consequently,

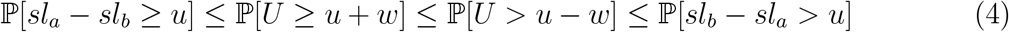

where the first inequality is from (3) and the third from (2). The inequalities are strict unless *U* does not have mass in (*u* − *w, u* + *w*]. Since *X_b_/Y_b_* and *X_a_/Y_a_* have the same distribution then *U* is sure to have positive probability in (−*w, w*). Thus with *u* = 0 and since ℙ[*sl_a_* − *sl_b_* ≥ 0] = ℙ[*sl_b_* − *sl_a_* ≤ 0] = 1 − ℙ[*sl_b_* − *sl_a_* > 0], then we have

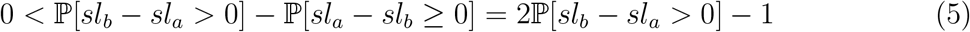

or ℙ[*sl_b_* − *sl_a_* > 0] > 1/2.

We have shown that under the alternative hypothesis that 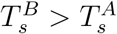 then we have that ℙ[*sl_b_* − *sl_a_* > 0] > 1/2. Under the null hypothesis that 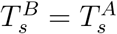 the distributions of *sl_b_* and *sl_a_* are the same. Thus if the actual normalized stem lengths were used for the two groups, the null and alternative hypotheses of interest imply the null and alternative hypotheses of the Mann-Whitney U test. However, the actual stem lengths are not known for the two groups; only estimates of these quantities from sequence data are available. This raises the question: Will the null and alternative hypotheses 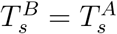 and 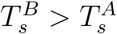 correspond to Mann-Whitney U test null and alternative hypotheses if estimated stem length distributions are used?

Assume that the number of sites is sufficiently large for each gene that asymptotic likelihood theory gives a good approximation to the sampling distributions of the stem lengths. That theory implies that 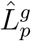 is approximately normal with mean 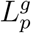. It follows from delta-method arguments (cf. §5.3.2 of Bickel and Doksum 2007) that 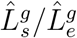 is approximately normal with mean 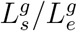 for any path *e* from LECA to a eukaryotic taxon. Because there are finitely many paths, for relatively large sequence lengths, the path, *e*\* say, corresponding to the median 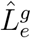 should coincide with the path corresponding the median 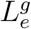. Thus 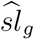 will be 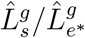 for the path *e*\* corresponding to the median 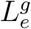. Since 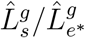 is approximately normal with mean 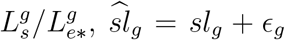 where ϵ_*g*_ has a normal distribution that is symmetric around 0. Consequently 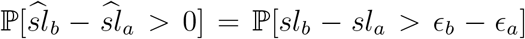. As a difference of independent, symmetric normals, *ϵ_b_* − *ϵ_a_* is symmetric normal. Denote the probability density function of the latter as *p*(*u*). Then

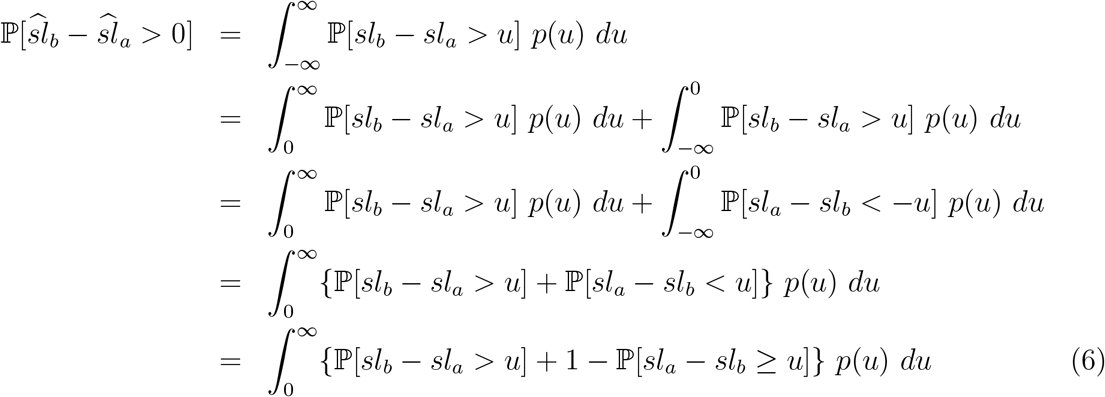

By (4), under the alternative hypothesis, ℙ[*sl_b_* − *sl_a_ > u*] − ℙ[*sl_a_* − *sl_b_* ≥ *u*] ≥ 0 with strict inequality in a neighbourhood of 0. Thus

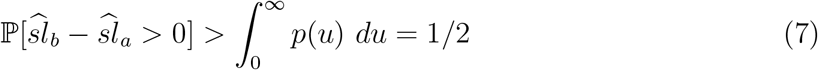

We see that the alternative hypothesis of interest corresponds to 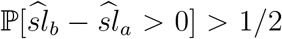 as required for the Mann-Whitney U test. However, the situation under the null is a little more problematic. Under the null hypothesis, ℙ[*sl_b_* − *sl_a_ > u*] − ℙ[*sl_a_* − *sl_b_* ≥ *u*] ≤ 0, so (6) gives that

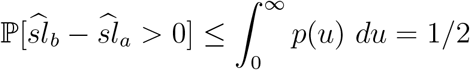

However, the Mann-Whitney U test requires that the distributions of 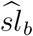 and 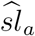 be the same. Although the distributions of the *sl_a_* and *sl_b_* are the same and the distributions of *ϵ_a_* and *ϵ_b_* are both symmetrically normal, their variances need not be comparable. These variances reflect precision of estimation and reasons that they might differ include that numbers of sites in alignments tend to differ substantially for one group versus the other. The null distribution used by the Mann-Whitney U test is not correct in such settings. Indeed, Kyusa (2000) shows that if the distributions for the two groups considered by the Mann-Whitney test are normal but with differing variances, the type I error of the test can be inflated. Nevertheless, Chung and Romano (2015) provide an alternative test that can be used under the null hypothesis, 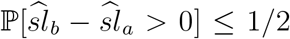 but 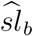 and 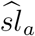 do not have the same distribution and we recommend use of this test as a safeguard. That being said, if the variability of estimation is comparable for the two groups, a Mann-Whitney U test should give reasonable results.

Much of the preceding discussion considers a case arising in both the analyses of Pittis and Gabaldón (2016a), and Vosseberg and colleagues (2020) where the times of origin associated with a stem length are constant for proteins within a group (for instance because they all derived from a mitochondrial symbiont or were all inherited from the archaeal host). Pittis and Gabaldón, and Vosseberg and colleagues, however, also compared groups made up of proteins of different bacterial origins and considered functional classes of proteins as groups. In these cases, proteins within a group are not expected to have a single time of origin (i.e., 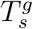 will vary within a group).

To allow for stem times that are not constant within groups, we assume a model in which 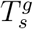 are independent across genes and independent of the rate variation processes 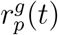. With the previous assumptions, the null hypothesis (that for *a* ∈ *A* and *b* ∈ *B*, 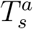 and 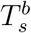 have the same distribution) implies that *sl_a_* has the same distribution as *sl_b_*. The alternative hypothesis of greatest interest is that there is no overlap in the 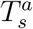 and 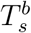 distributions: that 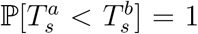. With this assumption, the arguments assuming fixed 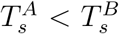, apply for the conditional distribution of 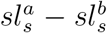, given 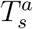 and 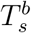. Averaging over 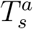 and 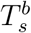 give that ℙ[*sl_b_* − *sl_a_* > 0] ≤ 1/2 under the null hypothesis and ℙ[*sl_b_* − *sl_a_* > 0] > 1/2 under the alternative hypothesis.

One difficulty with analyses when the 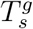 vary within groups is that we have no control over the alternative hypothesis. The desired alternative conclusion is that 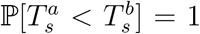 and we have argued above that such an alternative relationship leads to a Mann-Whitney U test null and alternative hypotheses for 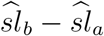. But suppose now that

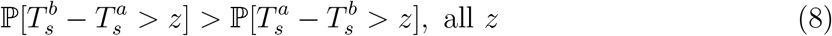

This condition is related to the hypothesis of interest but might not be very meaningful. For instance, if 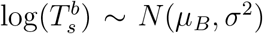 and 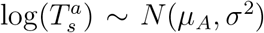 with *μ_B_ > μ_A_*, (8) holds but if *σ*^2^ is large then there is a substantial chance that any given 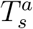 is larger than a given 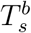. In other words, if substantial numbers of proteins in a given group *A* have an older origin than many of the proteins in group *B*, then what should we conclude from rejecting the null hypothesis that the group *A* distribution is shifted to be older than the group *B* distribution? More broadly, the rationale for grouping proteins together to test hypotheses about timings of origin is unclear if the age ranges across proteins in the groups heavily overlap and there is large variation within them.

We now show that (8) can lead to 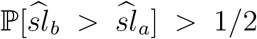. Assume for simplicity that ℙ[*r_p_*(*t*) = 1] = 1. Then Then the arguments leading to (4) apply with random 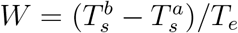 and exact equality:

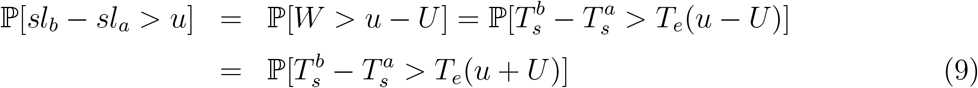

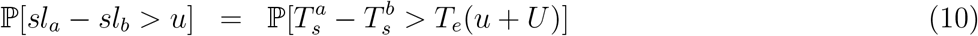

where the last equality in (9) and the equality in (10) follows from independence and the symmetric distribution of *U*. Letting *Z* = *T_e_*(*u* − *U*), then

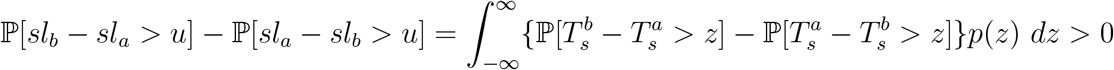

It follows as before that 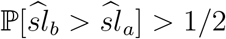. Since the Mann-Whitney U test or the Chung and Romano robust alternative are designed to detect 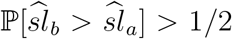 vs 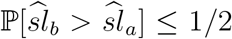, whatever the cause, rejection could correspond to less meaningful alternatives like those discussed above.

Another concern arises specifically for groups of genes that were laterally acquired during eukaryogenesis from a prokaryotic lineage. As discussed above and shown in Fig. 1, the actual stem-length time *T_s_*\* for these genes is less than the stem-length time *T_s_* for the observed tree. Throughout the preceding discussion we have assumed that 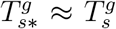. Suppose now that the two groups *A* and *B* have different prokaryotic origins and that 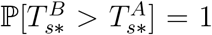. Will the Mann-Whitney U test be likely to reject in this case? Let 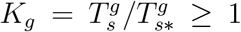. Recall that 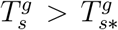 is expected because an immediate extant sister group to the actual prokaryotic transfer lineage is unlikely to be among the sampled taxa because of extinction. If the extinction processes are roughly the same for the two prokaryotic origin groups, then it is plausible that *K_g_* will have the same distribution for the two groups. We thus make the additional assumption that the *K_g_* have the same distribution for the two groups and are independent of the rate process below.

We now argue that the Mann-Whitney U test is indeed likely to reject when two such acquired groups of genes *A* and *B* have different single prokaryotic origins and origin times are well separated: 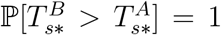. We condition on fixed 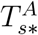 and 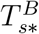 in what follows. Because the result below holds for all fixed 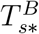 and 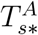, then averaging with respect to the distribution of 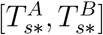 gives the result for random 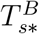 and 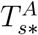.

Similarly as when *T_s*_* = *T_s_* was assumed, *sl_g_* = *X_g_/Y_g_* for *g* ∈ *A* and *sl_g_* = (*X_g_* + *V_g_*)/*Y_g_* for *g* ∈ *B*, where *Y_g_* is as in (1) but now

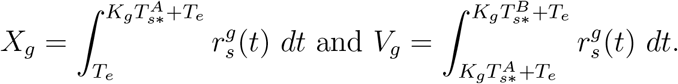

For *a* ∈ *A* and *b* ∈ *B*, observe first that, because the {*r_b_*(*t*)} and {*r_b_*(*t*)} processes are probabilistically equivalent, and because *K_a_* and *K_b_* have the same distributions, then *X_b_/Y_b_* and *X_a_/Y_a_* have the same distribution. Second, because *K_g_* ≥ 1, the smallest *V_b_/Y_b_* can be is then 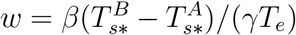. These two properties were what was used in the arguments leading to (2)–(5) and so those results hold in this setting. Here (5) gives the conclusion required for the Mann-Whitney U test, that ℙ[*sl_b_* − *sl_a_ >* 0] > 1/2 and (2) is the key inequality that can be used, exactly as before, to show (7), that, even with estimation, 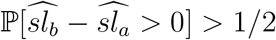.

Note that the above assumption that *K_g_* has the same distribution across groups is violated for some comparisons. For instance, comparisons of distributions from groups of acquired genes (where 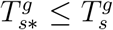 and, possibly, broad ranges of 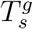 values within the group) with genes inherited from the asgard archaeon-eukaryote common ancestor or genes that originate by duplication (for which 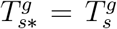 in both cases). In the simplest case of fixed 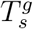 values for the genes in an acquired group, the inferred age of that group will be biased to be older than its true age in comparison with genes inherited from the asgard-eukaryote common ancestor or groups of duplicated genes.

We have shown that validity of the edge length ratio methods introduced by Pittis and Gabaldón (2016a), and extended by Vosseberg and colleagues (2020) do not require a molecular clock. They can be justified in much more general settings where substitution rates in a protein stochastically vary over the tree. Indeed, the only restrictions are that the stochastic substitution rate process is bounded away from 0 and infinity and that genes in groups of different origins (or functional classes) in a genome are all independently evolving according to this same process (i.e. the rate process for different genes are probabilistically equivalent). In terms of biological realism, it is this latter assumption that may not always hold. For example, it is well known that proteins may periodically experience episodes of rapid adaptive evolution due to acquisition of novel functions and/or loss of ancestral functions (Studer, Dessailly and Orengo, 2013). If this functional divergence differentially affected proteins within groups of different origins or functional classes, then stem length distributions of one group of proteins versus another will likely reflect this episodic shift in rates in one group, and cannot be used to test if they originated at different times. When applying these methods, it is therefore important to investigate evidence for systematic differences in the evolutionary dynamics of one group of proteins versus another.

It is also important to select edge lengths from phylogenies for different gene groups to be compared in the same manner to ensure that no biases are introduced. For example, when extending the approach to genes duplicated during eukaryogenesis, Vosseberg and colleagues (2020) were faced with deciding how to deal with the multiple possible edges or paths to LECA nodes created by duplications (Fig. 2). Their approach was to always select the minimum of all possible edge or path lengths for calculations of stem lengths or duplication lengths (all duplication lengths were calculated as 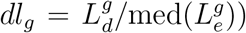). For two groups *D* and *S* of duplicated or acquired genes (for which stem lengths can include duplicated edges), the alternative hypothesis of greatest interest is *H_DS_*: ℙ[*T_di_* > *T_sj_*] = 1 for *d* ∈ *D* and *s* ∈ *S*, regardless of *i* and *j* ∈ {1, 2}.

**Figure 2:**
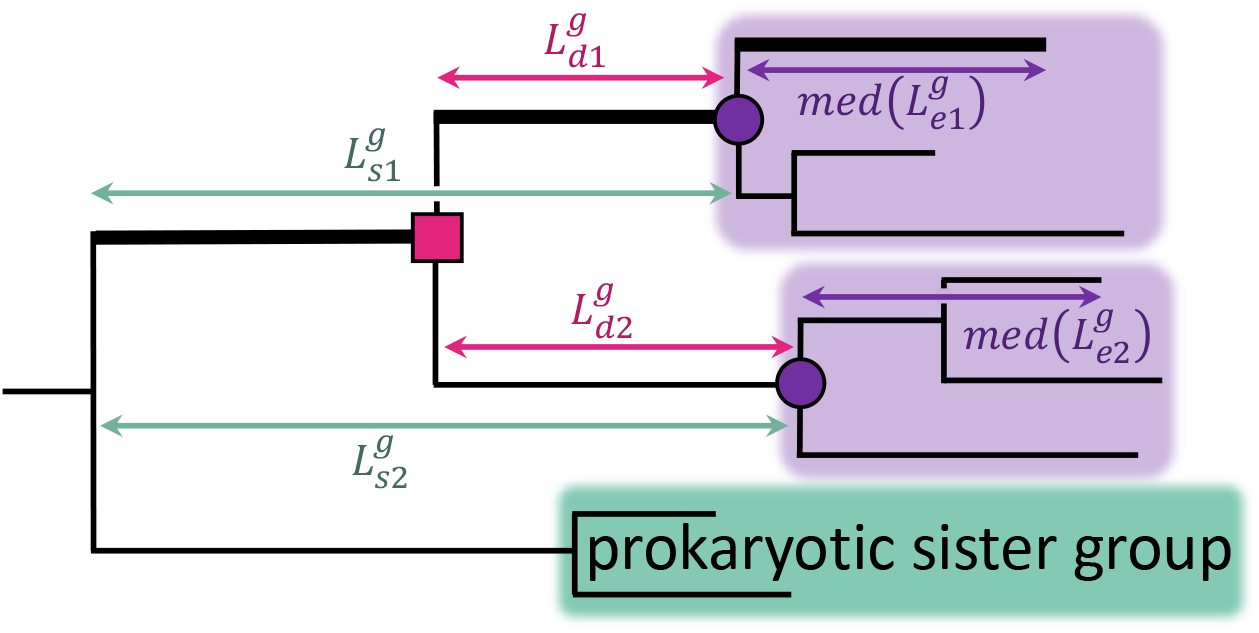
Example of the multiple possible stem paths and duplication edges in the phylogeny of a protein that has been duplicated during eukaryogenesis. This gene was acquired by the proto-eukaryote genome from a prokaryotic lineage (green box) and then duplicated (magenta box) prior to LECA (the LECA nodes of each duplicate are shown as purple circles). As a result there are two possible stem paths with lengths 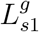 and 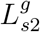 (green arrows), two possible duplication edges with lengths 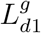 and 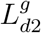 and two possible median paths within the eukaryote subtrees (purple boxes) with lengths 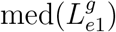 and 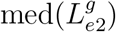. To calculate stem lengths and duplication lengths, Vosseberg and colleagues (2020) used the edges/paths with minimum values (minimums shown as thicker edges).

Assume that evolution is independent post duplication. Then, if all duplicated lengths are included in a comparison involving minima, the arguments above apply and the Mann-Whitney U test would be likely to reject when *H_DS_* holds. One possible motivation for using minima is that it could potentially alleviate bias due to the functional divergence phenomenon alluded to above because functional divergence is more likely to have occurred in the duplicate with the longer 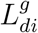 over *i* ∈ {1, 2}. Unfortunately, taking minimums leads to some loss of information and could lead to a bias which we illustrate assuming no functional divergence. Similar arguments as those above related to the median path lengths, imply that the Mann-Whitney U test would have substantial probability of rejection whenever ℙ[min(*T_d_*_1_, *T_d_*_2_) > min(*T_s_*_1_, *T_s_*_2_)] = 1. If *H_DS_* holds and evolution is independent, then this hypothesis will hold, so the approach should work. In less ideal alternative hypothesis scenarios where there is overlap in the distributions of *T_di_* and *T_sj_*, biases can occur. As an illustrative example, suppose that *S* is a stem-length group of acquired genes, that *T_e_* = 1.5 billion years ago, that *T_sj_* = 0.4 billion years and that, independently, *T_di_* ∼ *U* (0, 1). Then

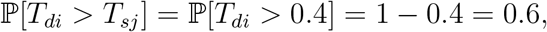

consistent with longer duplication lengths. But

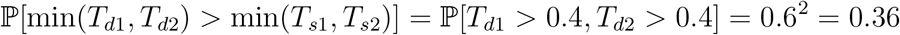

Consequently, based on the Mann-Whitney U test, one would conclude that the age of the duplicated group of genes *D* tended to be less than the acquired genes *S* when, in fact, 60% of the duplication group genes, *D*, duplicated prior to their acquisition.

Another complication arises in the comparison of duplication groups and stem-length groups. For duplication groups, duplication lengths are all minima. The stem-length groups, however, are usually a mix of stem lengths, some of which are chosen to be minima (Fig. 2) and some of which did not require a minimum (Fig. 1) (Vosseberg *et al.* 2020). In this case biases might arise even under the strong hypothesis, ℙ[min(*T_d_*_1_, *T_d_*_2_) > min(*T_s_*_1_*, T_s_*_2_)] = 1 for *d* ∈ *D* and *s* ∈ *S*, making it difficult to reject when *H_DS_* holds. Such problems can be averted by including all duplication and stem lengths rather than minima. There are potentially other ways of addressing functional divergence that may be less likely to introduce bias (e.g., identifying functionally divergent sites using methods reviewed in Studer, Dessailly and Orengo (2013) and removing them prior to analysis).

In summary, although there are a number of caveats, if the assumptions of the methods we have elaborated above are met by the data, these edge length ratio methods have the potential to provide important new insights into the roles of gene duplication and gene invention in different cellular systems and clarify the relative contributions of host, symbiont and lateral transfers to a lineage of interest.

## Acknowledgements

This research was supported a grant (Award ID: 735923LPI) awarded to A.J.R. and E.S. as part of the Moore-Simons Project on the Origin of the Eukaryotic Cell. E.S. and A.J.R. also acknowledge partial support for this work from Discovery grants awarded to them by the Natural Sciences and Engineering Research Council of Canada. A.J.R. acknowledges the Allan Wilson Centre in New Zealand for sabbatical support received in 2006 when some of these ideas were first discussed.

